# Ultra-accurate sequencing reveals an extreme transmission bottleneck in a deep-sea clam symbiosis

**DOI:** 10.64898/2026.06.29.735038

**Authors:** Cade Mirchandani, Evan Pepper-Tunick, Landen Gozashti, Shelbi Russell, Russ Corbett-Detig

## Abstract

Vertically transmitted symbionts experience progressive genome degradation driven by transmission bottlenecks each host generation that reduce genetic diversity and promote fixation of deleterious mutations. Direct estimates remain rare because inference requires scarce parent–offspring samples and sequencing sensitive enough to detect rare variants. Here, we investigate symbiont transmission bottlenecks in a vesicomyid clam by deeply sampling within-host endosymbiont genetic diversity using two ultra-accurate sequencing methods. Demographic modeling revealed an effective bottleneck size of approximately eight symbionts (95% CI: 1–17 genomes) per host generation. This estimate is sharply reduced relative to prior cytological estimates of bottleneck census size, with important implications for understanding the rate and dynamics of endosymbiont genome degradation.

## Introduction

Obligate endosymbionts experience progressive genome reduction driven by small effective population sizes, reduced recombination, and recurrent transmission bottlenecks that intensify genetic drift (1). Over time, this process drives the irreversible accumulation of deleterious mutations through Muller’s ratchet (2). The severity of genome erosion depends critically on the number of genetically distinct symbiont lineages transmitted from parent to offspring, termed the transmission bottleneck size (3).

Despite its importance in determining evolutionary outcomes, the transmission genetic bottleneck size has proven difficult to measure directly. Existing estimates typically rely on cell counts rather than genetically distinct lineages, and these may differ substantially: cell counts may not reflect genetically distinct founders and the tissue sampled may not capture the maximal bottleneck. In the vesicomyid Calyptogena okutanii, roughly 400 symbiont cells are transmitted per egg, each containing approximately 10 genome copies (4), but because the transmission bottleneck may precede embryo provisioning, these cytological counts may not reflect the number of genetically distinct founders. In the aphid-Buchnera system, approximately 1,800 symbiont cells are transmitted per egg (5), yet intergenerational symbiont allele frequency dynamics suggest within-host effective population sizes of only 10–20 genomes. Obtaining parent–offspring samples is impractical in many systems including deep-sea clams, and standard sequencing lacks the sensitivity to characterize within-host allele frequency variation.

Although strict vertical transmission is rare among marine symbioses (6), the vesicomyid *Calyptogena magnifica* and its sulfur-oxidizing symbiont Candidatus *Ruthia magnifica* exhibit predominantly vertical transmission (7), which has likely dominated for much of their estimated 70–80 million year obligate association, making this system particularly well-suited for studying endosymbiont bottleneck dynamics. Despite this ancient association, the symbiont genome (~1.2 Mb) remains larger and less degraded than terrestrial insect symbiont counterparts of comparable age, possibly due to occasional horizontal transmission and recombination (8), but the magnitude of the transmission bottleneck and its implications for genome degradation remain unknown. Here, we use two ultra-accurate sequencing methods to detect rare genetic variants within a single host individual and infer the effective transmission bottleneck size through demographic modeling.

## Results and Discussion

We applied two ultra-accurate sequencing methods to the same DNA extraction from the gill tissue of a single vesicomyid clam collected from the Galápagos Rift (0°48.2’N, 86°13.9’W, 2,461 m depth). Whole-genome alignment of the symbiont consensus assembly to the *Ca*. Ruthia magnifica reference genome (GCF_000015105.1) yielded 99.98% nucleotide identity (212 SNPs and 24 indels across 1.16 Mb) and 100% identity at the 16S rRNA gene. Because *Ca*. Ruthia magnifica is an obligate, species-specific symbiont of *C. magnifica (9, 10)*, we infer the host accordingly. From this sample, circle sequencing (11, 12) identified 289 variants while index sequencing (13) identified 668 variants (Fig 1A). In both datasets, variants were rare (Fig 1B), with allele frequencies consistent with a largely clonal endosymbiont population.

**Figure 1.**
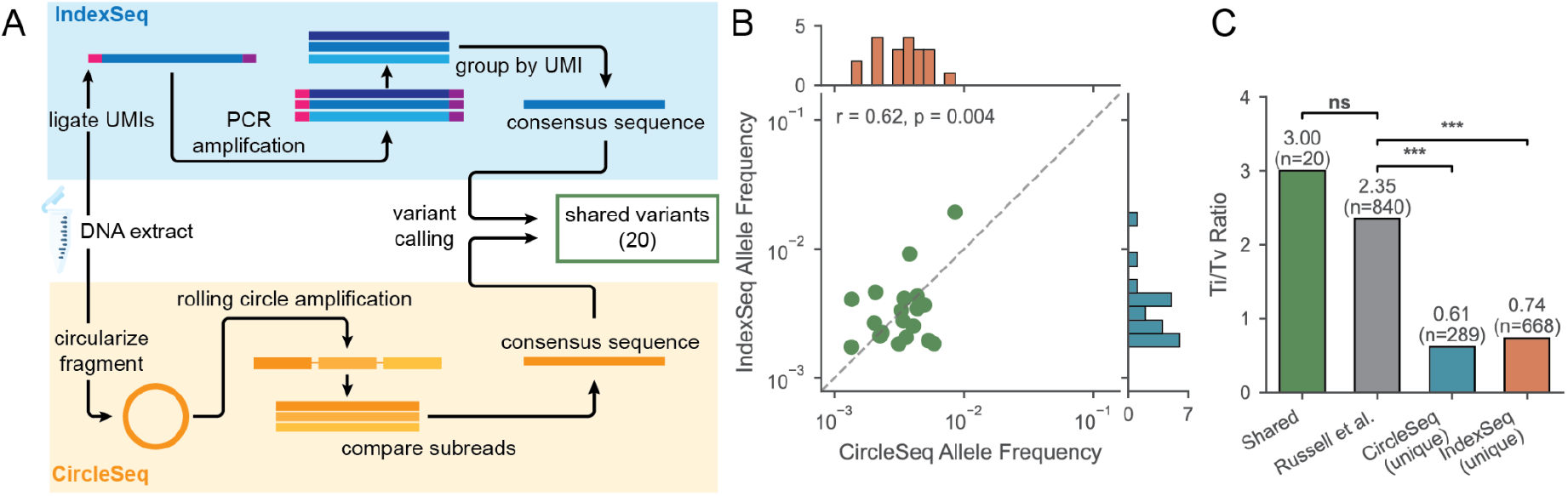
Variant validation. (A) Overview of sequencing approaches and variant calling pipeline. (B) Scatter plot showing allele frequency correlation between CircleSeq and IndexSeq at 20 shared variant positions, with marginal frequency distributions for each method. (C) Transition-to-transversion (Ti/Tv) ratios for shared variants, Russell et al. (2020) variants, and variants unique to each sequencing method. Chi-squared tests compare each category to Russell et al.; shared variants show no significant difference (χ^2^ = 0.05, p = 0.82), while variants unique to CircleSeq (χ^2^ = 92.35, p < 0.001) and IndexSeq (χ^2^ = 116.39, p < 0.001) differ significantly.

The significant overlap in variant calls and plausible biological properties of variants detected by each method indicates both methods identify a true biological signal. The two approaches detected largely distinct variant sets, with 20 variants shared between them, presumably due to extremely low allele frequencies and method-specific systematic errors. This overlap was significantly greater than expected by chance (permutation test, p < 0.001). Allele frequencies at shared positions were significantly positively correlated between methods (Pearson r = 0.62, p = 0.004) and ranged from 0.0014 to 0.0193 (Fig 1B), which suggests that for these overlapping variants, both methods detected a largely concordant biological signal. Finally, the transition-to-transversion (Ti/Tv) ratio of shared variants was comparable to that of polymorphic variants across a population of *C. Ruthia magnifica* sequenced with short-reads (Russell et al. 2020, p = 0.82, Fig 1C), whereas variants private to either sequencing method demonstrated substantially reduced Ti/Tv ratios (Fig 1C). Together, these results suggest that the shared variants represent genuine biological mutations suitable for demographic inference.

To infer the transmission bottleneck size from these shared variants, we performed demographic modeling of endosymbiont populations using a type of approximate Bayesian computation. We developed a discrete generation backward-in-time coalescent simulator to model symbiont genealogies under a two-phase demographic model of symbiont transmission (Fig 2A), and trained a random forest model on the resulting simulated allele frequency spectra to predict the bottleneck population size. The model predicted log_10_(Nb) with high accuracy (r = 0.80 on held-out simulations; Fig 2B), and predictions were largely independent of nuisance parameters population size (N), per-site mutation rate (μ), and stasis generations (g), though Nb weak residual correlation with μ (r = 0.24) because both parameters influence the number of detectable variants (Fig 2C). Applied to our 20 high confidence variants, the model estimated an effective bottleneck estimate of approximately 8 symbionts (95% CI: 1–17) per host generation (Fig 2D).

**Figure 2.**
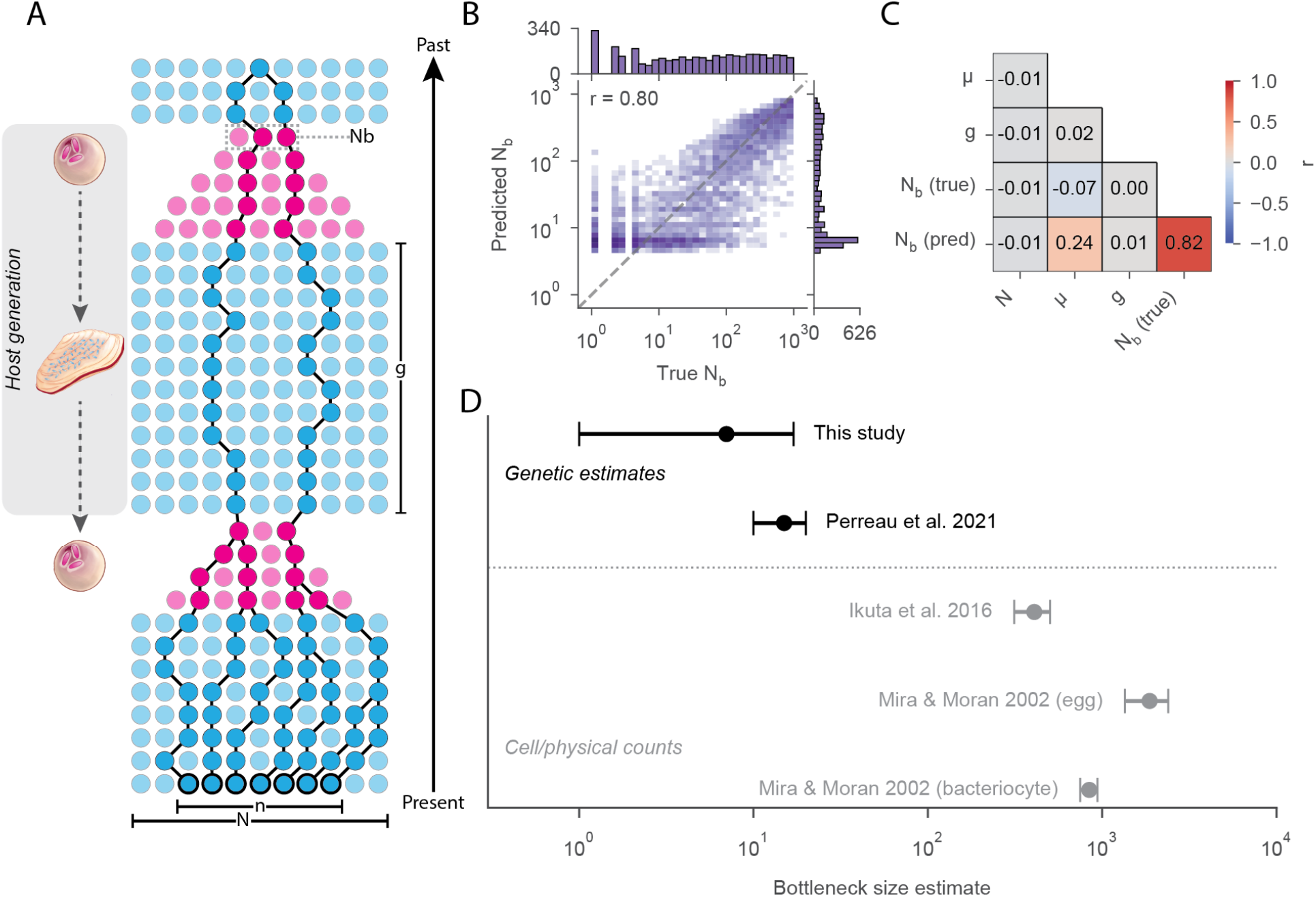
Bottleneck inference. (A) Two-phase demographic model. A host generation is shown on the left, with egg and adult clam cartoons indicating transmission. Circles represent symbiont lineages across generations; blue and pink denote stasis and bottleneck phases. Bold-outlined circles indicate sampled lineages (n). N, effective population size after host development; Nb, transmission bottleneck size; g, generations after initial expansion; per-site mutation rate (μ) not depicted. (B) Random forest performance on held-out simulations, with marginal distributions of true and predicted Nb. (C) Pairwise correlations among nuisance parameters (N, μ, g) and true/predicted Nb. (D) Bottleneck estimates across symbiont systems: genetic (black, effective genomes) vs. census (gray, microscopy).

Our estimated population bottleneck size is comparable to effective population size estimates in terrestrial insect endosymbionts, and substantially smaller than those derived from microscopy. Experimental evolution in aphid–Buchnera estimated within-host effective population sizes of only 10–20 (14), despite vastly different transmission mechanisms: vesicomyid symbionts are attached extracellularly to broadcast-spawned eggs (4), whereas Buchnera is transferred intracellularly from maternal bacteriocytes to developing embryos. Even under this severely restricted bottleneck, the *C. ruthia magnifica* genome remains substantially larger than terrestrial insect symbiont counterparts of comparable evolutionary age. This contrast lends further support to the inference that occasional horizontal transmission and recombination are more common in this clam population and act to maintain genome integrity in this system, counteracting the degenerative effects of drift even under extreme bottlenecks (8).

Genetic estimates of transmission bottlenecks complement existing census-based observations by indirectly inferring the number of genetically independent lineages transmitted each host generation. For example, in the related *C. okutanii*, approximately 400 symbiont cells are found on host egg surfaces (4), and in aphid–*Buchnera*, roughly 1,800 cells are transmitted per egg (5). Our estimate is orders of magnitude below these census counts, consistent with the expectation that effective and census population sizes diverge substantially in microbial systems (15). While census counts provide valuable insight into the physiology and ecology of symbiont transmission, the forces shaping genome erosion tend to be driven by effective population size. Therefore, genetic estimates across diverse host phylogenies, transmission mechanisms, and reproductive modes will be necessary for accurately parameterizing the evolutionary forces shaping endosymbiont genome trajectories.

## Methods

We called variants from two ultra-accurate sequencing methods applied to a single DNA extraction from *C. magnifica* gill tissue. Bottleneck size was inferred by training a random forest regressor on simulated allele frequency spectra generated under a discrete backward-in-time demographic model. Full details are in SI Appendix.

## Acknowledgements

We are grateful for the computational resources provided by the UCSC Genomics Institute. We thank Alexandra Lum for artwork and input on figure design. This work was supported by the National Institutes of Health grant R35GM128932 (to RCD), SLR was funded by National Institutes of Health grant R00GM135583.

## Declaration of Interests

R.C.-D. reports consulting for International Responder Systems. This consulting relationship had no role in the design, conduct, interpretation, or publication of this study. The authors declare no other competing interests.

## SI Appendix

### Sample Collection

Gill tissue was collected from a vesicomyid clam at the Galápagos Rift hydrothermal vent field (0°48.2’N, 86°13.9’W, ~2,461 m depth) on May 29, 1990 (Dive 2224) and stored at −80°C. The specimen was field-identified as a vesicomyid. To confirm species identity, we aligned a Pilon-polished (Walker et al. 2014) symbiont consensus assembly to the *Ca*. Ruthia magnifica reference genome (GCF_000015105.1, ASM1510v1) using minimap2 (Li 2018) with the asm5 preset. The alignment recovered 99.98% nucleotide identity across the full 1.16 Mb reference genome (212 SNPs, 10 insertions totaling 20 bp, and 14 deletions totaling 38 bp). We additionally extracted the 16S rRNA gene (reference positions 1,081,269–1,082,814) from both the reference and our assembly and compared them using BLASTn (Altschul et al. 1990), yielding 100% identity over 1,546 bp. These values fall within expected intraspecific variation and confirm the symbiont as *Ca*. Ruthia magnifica, consistent with the host being *Calyptogena magnifica*. Tissue was sterile-dissected or subsectioned from previously sterile-dissected gill tissue as described in Russell et al. (2020). DNA was extracted from gill tissue of a single individual for use in both sequencing approaches.

### Sequencing and Variant Calling

We used two complementary approaches to detect low-frequency variants from the same DNA extraction. For circle sequencing, DNA fragments are circularized and amplified via rolling circle amplification, generating tandem copies of each original molecule; we aligned these reads to the reference genome using BWA-MEM, realigned subsequences using Smith-Waterman alignment, and collapsed them into consensus sequences, masking any base with disagreement across subreads (Lou et al. 2013). For index sequencing, molecular barcodes are ligated to DNA fragments, allowing reads derived from the same original molecule to be identified after sequencing. Reads sharing a barcode are grouped into families and collapsed into single-strand consensus sequences (SSCS), in which each base is determined by majority agreement among family members. While the Duplex-Seq-Pipeline (Kennedy et al. 2014) can further combine complementary SSCS from opposite strands into duplex consensus sequences (DCS), duplex formation requires sufficient sequencing depth to recover both strands of each original molecule. Our target genome yielded insufficient strand pairing for reliable DCS generation, so we used SSCS for variant calling, which retains higher yield at a moderately higher error rate (~10^−4^) compared to DCS (~10^−6^; Lou et al. 2013).

We generated pileups from consensus sequences using samtools mpileup (Li et al. 2009) and called variants using a custom Python script (https://github.com/cademirch/symbiont_transmission). We applied identical filters to both methods. First, to identify error-prone genomic regions, we modeled the expected number of variants per 1 kb window as a Poisson distribution with rate λ = V × W / G, where V is the total variant count, W is the window size, and G is the genome size. We excluded windows in which the observed variant count exceeded the 99th percentile of this expectation, removing regions with anomalously high variant density indicative of systematic alignment or sequencing artifacts. We computed outlier windows independently for each method and excluded the union of both sets. Second, we required a minimum of 3 supporting consensus reads and a minimum allele frequency of 0.15%, approximately one order of magnitude above the per-base error rates for circle sequencing (~2–5 × 10^−5^) and single-strand consensus sequencing (~10^−4^; Lou et al. 2013). We calculated allele frequencies as alternate read counts divided by total depth. We defined shared variants as positions with the same alternate allele called independently by both sequencing methods. At matched consensus depths, the two methods produced comparable variant counts, and the higher number of method-specific variants in circle sequencing is consistent with its broader depth distribution (336–6,012×) compared to index sequencing (1,269–1,871×).

### Backward Time Symbiont Population Simulation

We developed a discrete-generations backward-in-time population simulator to model symbiont genealogies under an alternating-phase demographic model (https://github.com/cademirch/symbiont_transmission). The simulation tracks a sample of n lineages backward through time. The demographic model cycles through successive host generations, each consisting of two phases: a stasis phase where the symbiont population maintains effective size N for g generations, and a growth phase modeling post-transmission population expansion. In forward time, N_b_ symbionts colonize a new host and expand via binary fission to N over log_2_(N/N_b_) generations. In backward time, we model this as an exponential contraction where the effective population size halves each generation from N to N_b_, increasing the coalescent rate as lineages approach the transmission event. In each generation, we draw the expected number of coalescent events from a Poisson distribution with rate k(k−1)/2N, where k is the current number of active lineages. For each coalescent event, we choose two lineages uniformly at random, merge them into a single ancestral lineage, and record the branch lengths from each child to the new parent. Note that we explicitly do not allow more than two lineages to share a single common ancestor in one generation – i.e., consistent with binary cell division. So long as n << N, merges including three or more lineages are exceedingly rare. The simulation alternates between stasis and bottleneck phases, proceeding backward in time until all sampled lineages coalesce to a single common ancestor.

We place mutations on the resulting genealogy following a Poisson process. We aggregate branch lengths by the number of descendant samples: a mutation occurring on a branch with d descendants appears at frequency d/n in the sample. For each descendant class, we compute the expected number of mutations as the product of the total branch length in that class and the genome-wide mutation rate (per-site rate × genome size) μ, and draw the realized count from a Poisson distribution with this expectation. To model the observation process, we independently subsample each mutation through both sequencing methods. For each mutation at true sample frequency p = d/n, we draw a consensus depth D from the empirical depth distribution of each sequencing method and then draw the observed alternate read count from Binomial(D, p). While the hypergeometric distribution more exactly models sequencing as sampling without replacement, the probability of sampling an alternate read is sufficiently low that replacement has negligible effect on subsequent draws, and the Binomial provides an equivalent approximation. We consider a mutation detected by a given method if the alternate read count meets or exceeds the minimum threshold. This framework captures the joint effects of demographic history, mutational input, and the stochastic detection process of each sequencing method, allowing us to evaluate how well observed variant counts and allele frequency spectra match expectations under different demographic scenarios.

### Bottleneck Size Estimation

To estimate the transmission bottleneck size, we trained a random forest regressor on simulated allele frequency spectra generated under varying demographic scenarios. We sampled 50,000 parameter combinations using a Sobol quasi-random sequence to achieve uniform coverage of the four-dimensional parameter space: bottleneck size (N_ᵦ_) from 1 to 1,000 (log-uniform), effective population size (N) from 10^8^ to 10^10^ (log-uniform), per-site mutation rate (μ) from 10^−10^ to 10^−8^ per generation (log-uniform), and stasis generations (g) from 100 to 10,000 (uniform). For each parameter combination, we generated an observable allele frequency spectrum using the sequencing observation model described above, applying the same variant-calling thresholds used on empirical data and retaining only mutations detected by both methods. We binned allele frequencies into 10 log-spaced bins spanning 1.5 × 10^−3^ to 1, with the lower bound set to match our minimum detectable allele frequency of 0.15%. Bins were computed independently for each sequencing method, yielding a 20-dimensional feature vector per simulation.

We trained a random forest regressor (scikit-learn; (Pedregosa et al. 2012)) to predict log_10_(N_ᵦ_) from these feature vectors, holding out 20% of samples as a test set. We selected hyperparameters via randomized search over 100 iterations of 5-fold cross-validation, minimizing mean squared error, yielding a forest of 492 trees with maximum depth 10, minimum samples per split of 9, and 30% of features considered at each split. We evaluated model performance on the held-out test set by R^2^, root mean squared error (RMSE), and mean absolute error (MAE), and confirmed generalization with 5-fold cross-validated R^2^ on the training set. We applied the trained model to our empirical shared variants and estimated prediction uncertainty from the distribution of predictions across individual trees in the forest, reporting the mean as the point estimate and the 2.5th and 97.5th percentiles as a 95% confidence interval.

## References

1. J. J. Wernegreen, Genome evolution in bacterial endosymbionts of insects. Nat. Rev. Genet. 3, 850–861 (2002).

2. M. E. Pettersson, O. G. Berg, Muller’s ratchet in symbiont populations. Genetica 130, 199–211 (2007).

3. M. Kaltenpoth, W. Goettler, S. Koehler, E. Strohm, Life cycle and population dynamics of a protective insect symbiont reveal severe bottlenecks during vertical transmission. Evol. Ecol. 24, 463–477 (2010).

4. T. Ikuta, et al., Surfing the vegetal pole in a small population: extracellular vertical transmission of an “intracellular” deep-sea clam symbiont. R Soc Open Sci 3, 160130 (2016).

5. A. Mira, N. A. Moran, Estimating population size and transmission bottlenecks in maternally transmitted endosymbiotic bacteria. Microb. Ecol. 44, 137–143 (2002).

6. S. L. Russell, Transmission mode is associated with environment type and taxa across bacteria-eukaryote symbioses: a systematic review and meta-analysis. FEMS Microbiol. Lett. 366 (2019).

7. S. C. Cary, S. J. Giovannoni, Transovarial inheritance of endosymbiotic bacteria in clams inhabiting deep-sea hydrothermal vents and cold seeps. Proc. Natl. Acad. Sci. U. S. A. 90, 5695–5699 (1993).

8. S. L. Russell, et al., Horizontal transmission and recombination maintain forever young bacterial symbiont genomes. PLoS Genet. 16, e1008935 (2020).

9. I. L. G. Newton, et al., The Calyptogena magnifica chemoautotrophic symbiont genome. Science 315, 998–1000 (2007).

10. F. J. Stewart, C. R. Young, C. M. Cavanaugh, Lateral symbiont acquisition in a maternally transmitted chemosynthetic clam endosymbiosis. Mol. Biol. Evol. 25, 673–687 (2008).

11. G. A. Penunuri, E. A. Pepper-Tunick, J. McBroome, R. Corbett-Detig, S. Russell, EMS mutation and SNP detection in intracellular Wolbachia genomes. bioRxiv (2026).

12. D. I. Lou, et al., High-throughput DNA sequencing errors are reduced by orders of magnitude using circle sequencing. Proc. Natl. Acad. Sci. U. S. A. 110, 19872–19877 (2013).

13. S. R. Kennedy, et al., Detecting ultralow-frequency mutations by Duplex Sequencing. Nat. Protoc. 9, 2586–2606 (2014).

14. J. Perreau, B. Zhang, G. P. Maeda, M. Kirkpatrick, N. A. Moran, Strong within-host selection in a maternally inherited obligate symbiont: Buchnera and aphids. Proc. Natl. Acad. Sci. U. S. A. 118 (2021).

15. L.-M. Bobay, H. Ochman, Factors driving effective population size and pan-genome evolution in bacteria. BMC Evol. Biol. 18, 153 (2018).

